# Increased early sodium current causes familial atrial fibrillation and dampens effect of flecainide

**DOI:** 10.1101/2022.01.18.476646

**Authors:** M O’Reilly, LC Sommerfeld, C O’Shea, S Broadway-Stringer, S Andaleeb, JS Reyat, SN Kabir, D Stastny, A Malinova, D Delbue, L Fortmueller, K Gehmlich, D Pavlovic, BV Skryabin, AP Holmes, P Kirchhof, L Fabritz

## Abstract

**Aims:** Atrial fibrillation (AF) is the most common cardiac arrhythmia. Pathogenic variants in genes encoding ion channels are associated with familial AF. The point mutation M1875T in the *SCN5A* gene, which encodes the α-subunit of the cardiac sodium channel Na_v_1.5, has been associated with increased atrial excitability and familial AF.

**Methods:** We designed a new murine model carrying the *Scn5a*-M1875T mutation enabling us to study the effects of the Na_v_1.5 mutation in detail *in vivo* and *in vitro* using patch clamp and microelectrode recording of atrial cardiomyocytes, optical mapping, ECG, echocardiography, gravimetry, histology and biochemistry.

**Results:** Atrial cardiomyocytes from newly generated adult *Scn5a*-M1875T^+/-^ mice showed a selective increase in the early (peak) cardiac sodium current, larger action potential amplitude and a faster peak upstroke velocity. Conduction slowing caused by the sodium channel blocker flecainide was less pronounced in *Scn5a*-M1875T^+/-^ compared to wildtype atria. Overt hypertrophy or heart failure in *Scn5a*-M1875T^+/-^ mice could be excluded.

**Conclusion:** The *Scn5a*-M1875T point mutation causes gain-of-function of the cardiac sodium channel. Our results suggest increased atrial peak sodium current as a potential trigger for increased atrial excitability and thus AF.

**What’s new:**

- The point mutation M1875T in the C-terminal domain of the cardiac sodium channel Na_v_1.5 causes an increase in early peak sodium current in left atria.
- The observed changes induced by this point mutation suggest an increase in peak sodium current as a cause of familial atrial fibrillation (AF).
- Our findings provide a possible explanation for the variable effectiveness of sodium channel blockers in patients with AF. Carriers of such sodium channel gain-of-function mutations may benefit more from tailored treatments.

**Graphical abstract:** 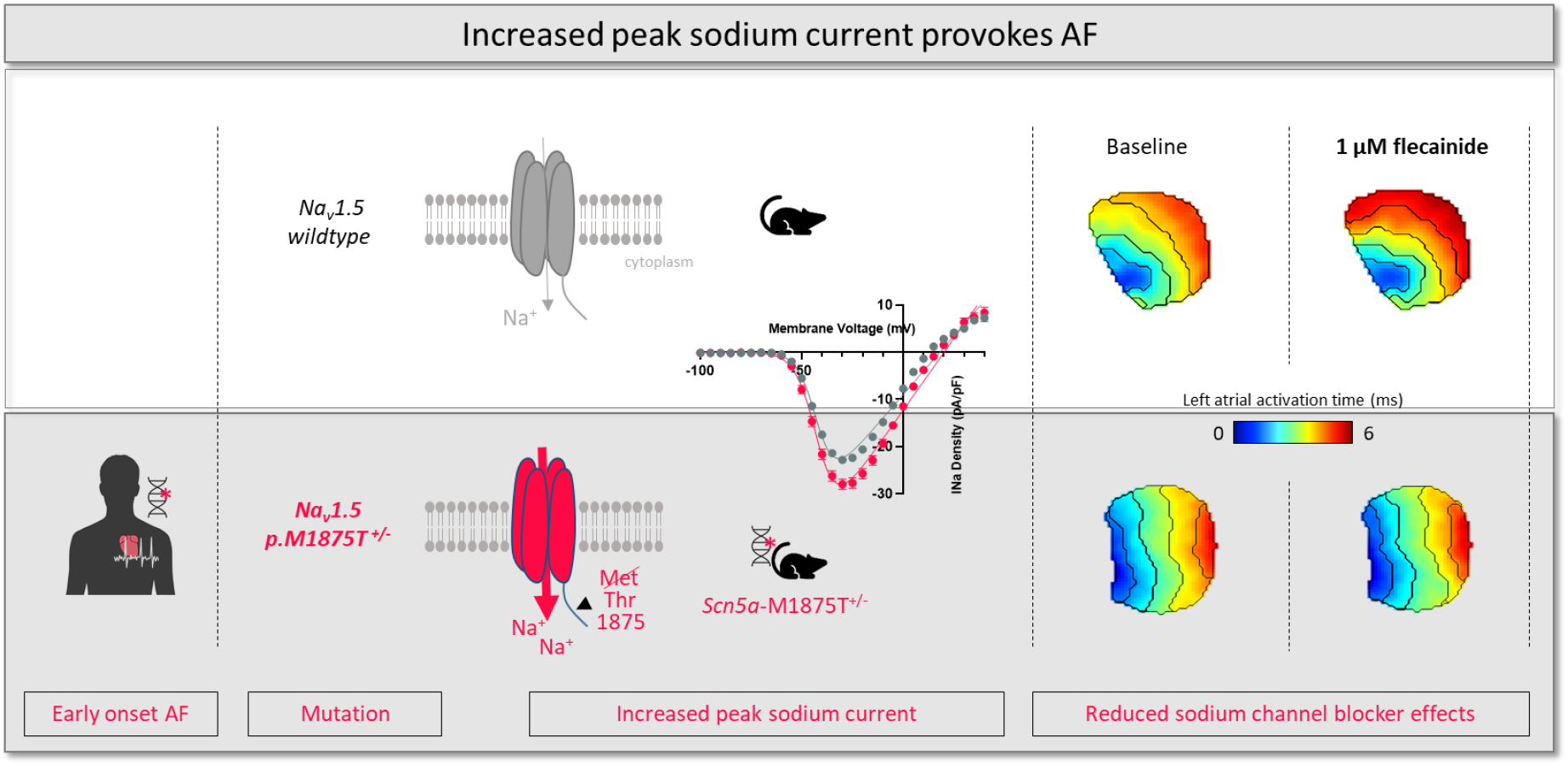

## 1 Introduction

Atrial fibrillation (AF), the most common cardiac arrhythmia, is characterised by episodes of irregular and uncoordinated atrial electrical activity. It is associated with ischaemic stroke, cardiovascular death and frequent hospitalisations ^1^. A range of different common factors, such as heart failure, diabetes and increased formation of fibrosis, can damage the atria, contributing to AF ^2, 3^. These factors interact with a pre-existent, potentially inherited substrate to result in AF. Inherited forms are characterised by early onset of the condition. Pathogenic variants in several genes associated with cardiomyopathies have been identified in familial AF, including variants in sarcomeric and cell-cell contact genes, and others ^4^. Within the group of ion channel genes, variants leading to dysfunction of the cardiac sodium channel are associated with familial AF ^4, 5^, both via variants in the genes coding for the channel or through upstream mechanisms altering the expression of sodium channel genes ^6, 7^.

The cardiac voltage-gated sodium channel (Na_v_1.5) facilitates movement of sodium (Na^+^) ions across the cardiomyocyte membrane and hence elicits the cardiac Na^+^ current (I_Na_). I_Na_ is vital for electrical excitation preceding mechanical contraction of the myocardium, as transient influx of Na^+^ triggers the fast upstroke phase of the cardiac action potential and thus depolarisation of the cardiomyocyte membrane. The Na_v_1.5 channel is composed of both α- and ß-subunits. The pore-forming α-subunit is encoded by the *SCN5A* gene. Variants in *SCN5A* have been linked to several cardiac conditions, including AF ^8–10^. *SCN5A* genetic variants reported show various underlying mechanisms mainly linked to channel dysfunction, defective channel trafficking or protein complex formation ^11, 12^.

A missense *SCN5A* point mutation, Met1875Thr (M1875T), located in the C-terminus of the channel protein, was linked to autosomal dominant familial AF that spanned three generations^13^. Atrial ectopy was evident in mutation carriers in adolescence, and persistent AF occurred as early as 27 years of age. Analyses in the human cell line HEK293 heterologous expression system ^13^ suggested an enhanced function of the mutated Na_v_1.5 channel.

To investigate the impact of the M1875T mutation in a more physiological setting, we generated and characterised a novel knock-in murine model (*Scn5a*-M1875T^+/-^). Mice that were heterozygous for the M1875T mutation were viable and studied herein. We investigated these mice from the whole organ *in vivo* to the level of the single cell *ex vivo*.

## 2 Methods

### 2.1 Generation and sequencing of the *Scn5a*-M1875T murine model

Mice heterozygous for the knock-in mutation M1875T in the *Scn5a* gene (*Scn5a*-M1875T^+/-^) were generated by T-C point-mutating exon 28 of the cardiac sodium channel *SCN5A* gene using pSCN5a_targ3 targeting vector and CRISPR/Cas9 system in murine embryonic stem (ES) cells (Figure 1a). Mutation-harboring ES cells were characterised using Southern blot analysis and sequenced to exclude genomic rearrangements (Figure 1b) ^14, 15^, and injected into B6D2F1 mouse blastocysts. Sequencing analysis of the mutation-containing region using DNA from adult wildtype (WT) and heterozygous *Scn5a*-M1875T^+/-^ mice on C57Bl/6J x 129sv hybrid genetic background are shown in Figure 1c. Detailed steps of the generation are further explained in the supplement.

**Figure 1.**
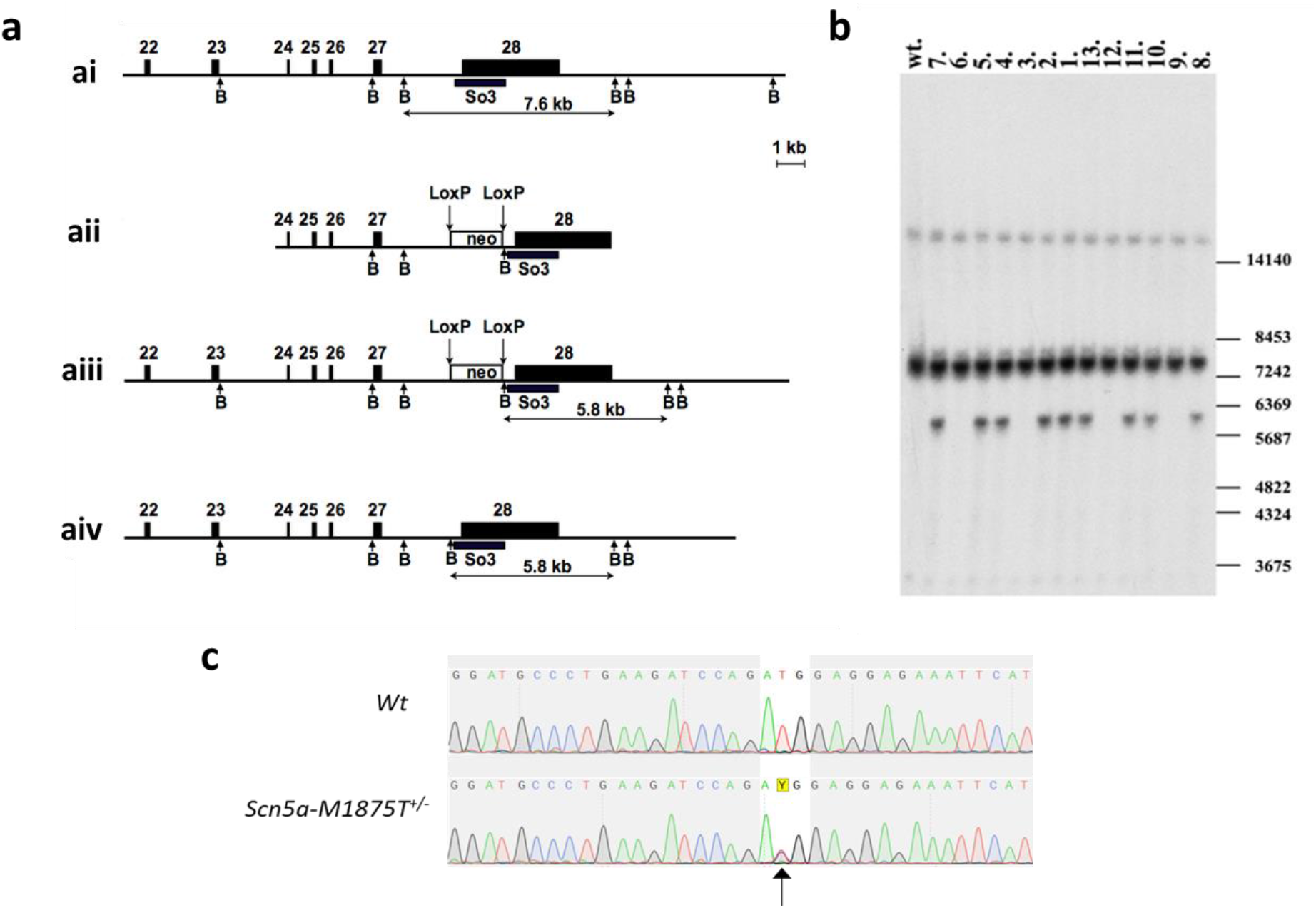
Targeting of exon 28 of the mouse *Scn5a* gene in order to introduce the M1875T coding mutation, and sequencing confirmation of its presence. **a)** The intronic and intergenic regions are shown as lines, exons are shown as filled boxes. The empty box corresponds to the neomycin resistance cassette (neo) flanked by the LoxP sites (vertical arrows). Exon numeration is shown above. The arrows below are corresponding to BamHI restriction endonuclease sites (B). The black box corresponds to Southern probe sequences (So3). The expected sizes of restriction DNA fragments are indicated below in kb. **ai)** Wildtype (WT) locus. **aii)** Targeted vector structure (without negative selection marker and plasmid backbone). **aiii)** Genomic locus after the homologous recombination. The neomycin cassette is present in intron 27 and flanked by two LoxP sites. **aiv)** Genomic locus after the CRE-mediated neo cassette deletion. **b)** Southern blot analysis of DNA isolated from mouse tail biopsy of the F1 offspring (1-13) and hybridized with the So3 probe. With help of the BamHI enzymatic digestion, we detect the WT allele 7.6 kb and targeted allele 5.8 kb. DNA samples 1, 2, 4, 5, 7, 8, 10, 11, and 13 contain correctly targeted *Scn5a* gene (Scn5a-M1875T^+/-^). Positions of the size marker (in bp) are shown on the right. The WT control animal is labeled “wt”. **c)** Sequencing analysis of the *Scn5a* gene region containing the mutation site in back-crossed adult mice (on pure genetic background). The T-C mutation on one allele causing the methionine-threonine exchange at position 1875 (1877) is indicated by an arrow.

Methionine at position 1875 of the human Na_v_1.5 protein sequence corresponds to position 1877 of the murine sequence. The latter is therefore point-mutated in this model. However, to underpin the bedside-to-bench nature of this investigation, we use the human annotation and refer to it as “Scn5a-M187**5**T” throughout this manuscript.

Mice were bred on an FVB or 129/sv genetic background and housed in individually ventilated cages with sex-matched littermates (2-5 mice/cage), under standard conditions: 12 hours light/dark circle, 22°C and 55% humidity. Food and water were available *ad libitum*. The health status of mice used in the study was monitored daily and prior to experiments.

Functional experiments were conducted on hearts of male and female young adult mice of pure background (8-20 weeks), heterozygous for the knock-in mutation M1875T in the *Scn5a* gene (*Scn5a*-M1875T^+/-^) and their WT littermates. Mice of both sexes and background were used evenly and analysed jointly, to make the data more widely applicable. Our investigations focus on the left atrium (LA). This is mostly due to practical reasons, e.g. less spontaneous activity in the left compared to the right atrium and therefore the possibility to pace at a wider range of cycle lengths or better reproducibility of the LA in the parasternal view.

### 2.2 Study approval

All procedures were performed in compliance with the guidelines from Directive 2010/63/EU of the European Parliament on the protection of animals used for scientific purposes and conducted in accordance with rules and regulations for experiments with animals and approved by the UK Home Office (PPL number 30/2967) and by the institutional review board of University of Birmingham.

### 2.3 ECG recordings *in vivo*

Non-invasive electrocardiograms (ECG) were recorded in conscious young adult mice (8-19 weeks) using a tunnel system for gentle restraint (ecgTunnel, EMKA Technologies, Paris, France) ^16^. ECG recordings were analysed using ECGauto software (EMKA Technologies, Paris, France). ECGs were also recorded in isoflurane-sedated mice during echocardiography as below.

### 2.4 Echocardiography *in vivo*

Echocardiography was performed in sedated mice (0.5-2% isoflurane, supplemented with 100% O_2_) using Vevo® 2100 system (VisualSonics, Amsterdam, Netherlands) as reported previously ^17^. Heart rate was maintained at 450 ± 70 bpm. Left atria (LA) were visualised in the parasternal long axis view in the plane of the aortic root. LA area and diameter were measured during pre-atrial contraction, using the P-wave of the limb ECG trace as a guide.

### 2.5 Histological analysis

Hearts were fixed in formalin and paraffin-embedded tissues were cut into slices of 4 μm. Sections were dewaxed, stained with hematoxylin and eosin for overviews and subsequently dehydrated, embedded and imaged on a NanoZoomer 2.0-HAT (Hamamatsu).

Slides used for quantitative analysis were cooked in citrate buffer for antigen retrieval. Autofluorescence was quenched with a 0.25% Sudan black solution for 30 min and samples were blocked with 2% BSA/2.2% Glycine for 1 h at room temperature (RT). Wheat germ agglutinin (WGA) with Alexa Fluor™ 488 Conjugate (1:200, W11261, Invitrogen) was applied for 2 h at RT.

Images of WGA-stained cardiac tissue were obtained with a confocal microscope equipped with an Aurox Clarity (Aurox Ltd.) spinning disc unit and a 20x EC Plan-Neofluar objective (420353-9900-000, Zeiss, NA=0.5) run with Aurox Visionary (Aurox Ltd.) software.

Quantitative analysis of WGA-stained area and cardiac cell diameters was carried out using a published ImageJ plugin for atrial histological analysis (JavaCyte^18^), with minor adjustments as outlined in supplementary methods.

### 2.6 Atrial cardiomyocyte isolation

Murine hearts were excised under deep terminal anaesthesia (4% isoflurane inhalation in O_2_, 1.5 L/min) and perfused at 4 mL.min^-1^ at 37°C on a vertical Langendorff apparatus with the following solutions, equilibrated with 100% O_2_: (i) HEPES-buffered, Ca^2+^-free, modified Tyrode’s solution containing in mM: NaCl 145, KCl 5.4, MgSO_4_ 0.83, Na_2_HPO_4_ 0.33, HEPES 5, and glucose 11 (pH 7.4, NaOH) x 5 min; (ii) Tyrode’s enzyme solution containing 640 μg/mL collagenase type II (270 U/mg), 600 μg/ml collagenase type IV (270 U/mg) and 50 μg/mL protease (Worthington, Lakewood, NJ), 20 mM taurine and 3 μM CaCl_2_ × 8-12 min. The heart was removed from the Langendorff setup and perfused with 5 mL of modified Kraftbruhe (KB) solution containing in mM: DL-potassium aspartate 10, L-potassium glutamate 100, KCl 25, KH_2_PO_4_ 10, MgSO_4_ 2, taurine 20, creatine 5, EGTA 0.5, HEPES 5, 0.1% BSA, and glucose 20 (pH 7.2, KOH).

The LA was dissected free and cardiomyocytes were dissociated gently with fire-polished glass pipettes (2 to 1 mm diameter in sequence). Cells were re-suspended in 2 mL KB buffer and Ca^2+^ was gradually reintroduced to the cell suspension incrementally over a period of 2 hours to reach a final concentration of 1 mM. All experiments were performed within 8 hours of isolation.

### 2.7 Whole-cell patch clamp electrophysiology of isolated atrial cardiomyocytes

Dissociated murine LA cardiomyocytes were plated on, and allowed to adhere to, laminin-coated coverslips (10 mm diameter) for at least 20 minutes. Coverslips were transferred to a recording chamber and were continually superfused at 3 mL.min^-1^, with a low Na^+^ external solution containing in mM; NaCl 10, KCl 4.5, C_5_H_14_CINO 130, CaCl_2_ 1, MgCl_2_ 1.2, HEPES 10 and glucose 10 (pH 7.4 with CsOH). To block L-type Ca^2+^ currents, 2 mM NiCl_2_ was added to the superfusate. Experiments were performed at 22 ± 0.5°C. Whole-cell patch clamp recordings were obtained in voltage-clamp mode using borosilicate glass pipettes (tip resistances 1.5-3 MΩ).

For Na^+^ current recordings, the pipette solution contained in mM: CsCl 115, NaCl 5, EGTA 10, HEPES 10, MgATP 5, TEACl 20 and MgCl_2_ 0.5 (pH 7.2, KOH). Voltage-dependent Na^+^ currents were evoked by 5 mV step depolarisations (100 ms) from a holding potential of −100 mV to test potentials ranging from −95 mV to +40 mV. Cells were excluded from analysis if there was no reversal of the sodium current by +40mV. To investigate Na_v_1.5 voltage-dependent inactivation kinetics, cells were subject to 500 ms pre-pulses ranging from −120 mV to −40 mV, followed by a 100 ms step to −30 mV. For Na_v_1.5 time-dependent recovery kinetics, a standard two pulse protocol was used (−120 mV to −30 mV, 20 ms), with the time between the two pulses incrementally varying between 5 and 950 ms.

All recordings and analysis protocols were performed using an Axopatch 200B amplifier (Molecular Devices, USA) and digitized at 50 kHz using a CED micro1401 driven by Signal v6 software (Cambridge Electronic Design, Cambridge, UK). Series resistance was compensated, ranging between 60-100% for all cells. Experiments were terminated if series resistance abruptly changed or was above 10 MΩ.

### 2.8 Atrial microelectrode recordings

As previously described ^19, 20^, following isolation the LA was immediately transferred into a dissecting chamber and continuously superfused at 10 mL.min^-1^ with a bicarbonate buffered Krebs-Henseleit (KH) solution containing in mM: NaCl 118; NaHCO_3_ 24.88; KH_2_PO_4_ 1.18; Glucose 11; MgSO_4_ 0.83; CaCl_2_ 1.8; KCl 3.52, equilibrated with 95% O_2_/5% CO_2_, 36-37°C, pH 7.4. Micro-dissection and pinning out of the LA was performed using a dissection microscope (Stemi SV 11, Zeiss, Germany). The LA was paced at 1–10 Hz via bipolar platinum electrodes. Action potentials (APs) were recorded from freely contracting LA using custom made glass floating microelectrodes containing 3 M KCl, (resistance 15-30 MΩ). Voltage signals were amplified and digitised at 20 kHz and were unfiltered (Axoclamp 2B; Molecular Devices, California, USA; Spike2 software Cambridge Electronic Design, Cambridge, UK). Measured parameters included the resting membrane potential (RMP), action potential amplitude (APA), peak depolarisation rate (dV/dt) and action potential duration (APD) at 30-90% repolarisation. APs were only analysed following sufficient rate adaptation achieved after at least 50 stimulated APs at each frequency.

### 2.9 Optical mapping of left atria

Optical mapping of the LA was conducted as previously described ^21, 22^. Isolated whole hearts were loaded on to a vertical Langendorff apparatus and perfused with a standard KH solution. Hearts were perfused at 4 mL.min^-1^ (equilibrated with 95%O_2_/5%CO_2_ and heated to 36-37°C, pH 7.4). Hearts were loaded with 25 μL of voltage sensitive dye Di-4-ANEPPS at a concentration of 5 mg/mL, diluted in 1 mL of KH solution and delivered via bolus port injection over 3-5 minutes. The LA was then isolated and pinned in a superfusion chamber containing 37°C KH solution for transfer to the optical mapping setup, anterior surface facing up.

In the optical mapping system, atria were superfused with KH solution (95%O_2_/5%CO_2_, 36-37°C) containing contraction uncoupler Blebbistatin (35 μM). For imaging, atria were illuminated by two dual LEDs at 530 nm. A 630 nm long-pass filter was used to sperate emitted fluorescence, imaged using an ORCA flash 4.0 CMOS camera (Hamamatsu, Japan). Images were acquired at a framerate 0.987 kHz and pixel size of 71 μm/pixel^2^. Atria were paced using bipolar platinum electrodes delivering 2 ms pulses at twice diastolic threshold (minimum voltage required to elicit APs).

One-minute baseline recording was taken following a 10 minute equilibration period to ensure contraction uncoupling and temperature re-stabilisation. During imaging, atria were initially paced at 330 ms pacing cycle length (PCL). A ‘ramp’ pacing protocol was then initiated, in which the atria were paced at 120 ms PCL for 100 stimuli and then PCL was reduced from 120 ms to 80 ms in 10 ms intervals every 20 stimuli. After taking baseline recordings, LED illumination was switched off and the superfusion solution replaced with an identical solution containing flecainide at a concentration of 1 μM and then 5 μM (or control solution without flecainide for time control experiments). Subsequent recordings were then made as described above after 20 minutes superfusion with 1 μM flecainide solution, and then further 15 minutes with 5 μM flecainide solution. Atria were paced at 330 ms PCL continuously in dark conditions between recordings.

From these recordings, APD and conduction velocity (CV) were mapped across the LA using ElectroMap software ^22^. Atria were removed from analysis at a given PCL if loss of 1:1 capture ratio with pacing stimuli (i.e. missed beats) was observed.

### 2.10 Statistics

For all murine experiments presented herein, experimenters were blinded to the genotype of the littermate pairs during data collection and analysis. Student t-tests were used for singular comparisons for normally distributed data. A hierarchical nested t-test (Figure 2) and Mann-Whitney test (Table S3) were used where appropriate. Multiple comparisons were made using 2-way ANOVA with Bonferroni’s post-hoc tests. For current-voltage graphs (Figure 3b), a Boltzmann curve was fit to the data, using the modified Boltzmann equation: I_Na_ = G_max_(V_m_ – V_rev_)/(1+Exp[(V_0.5_-V_m_)/k]), where I_Na_ is the current density at an equivalent test potential (V_m_), G_max_ is the peak conductance (nS), Vrev is the reverse potential, V_0.5_ is the membrane potential at 50% current activation, and k is the slope constant. All graphical representations display individual measurements. Means are quoted and shown in Figures ± SEM unless stated otherwise. Level of statistical significance is shown in Figures as follows: *p<0.05; **p<0.01; ***p<0.001; ****p<0.0001. Statistics and Figures were created using Prism 8 (GraphPad Software, San Diego, California).

**Figure 2.**
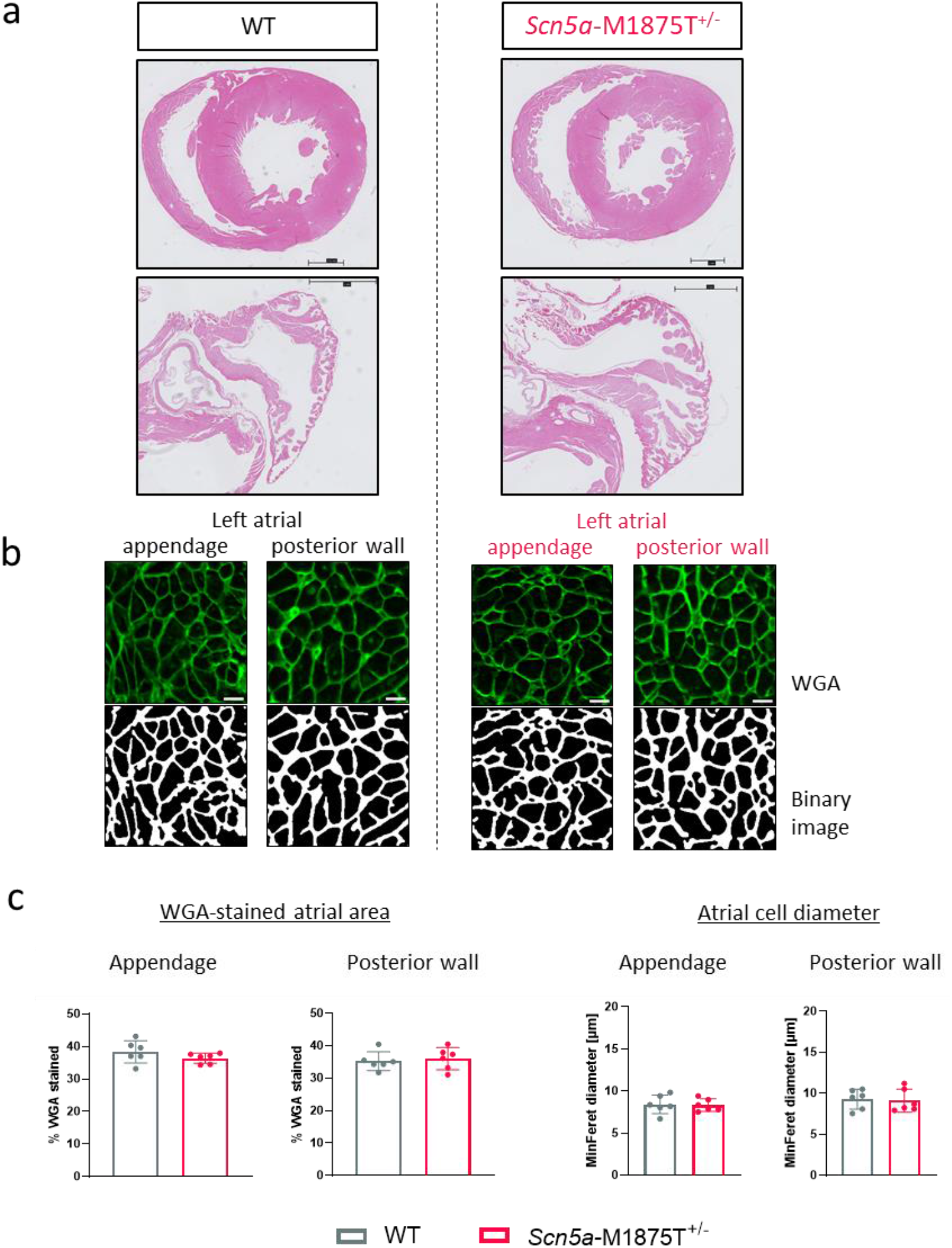
Unaltered atrial amount of extracellular matrix content and myocyte diameter. **a)** Hematoxylin and eosin-stained sections of wildtype (WT) and *Scn5a*-M1875T^+/-^ litter pair hearts (ventricles, upper panel; left atria, lower panel). Scale bars represent 1 mm. There were no obvious differences between genotypes. **b)** Exemplary immunofluorescence images from atrial regions of interest (ROI) of wheat germ agglutinin (WGA, green) staining and corresponding binary images for quantification in left atrial (LA) appendage and LA posterior wall from WT and *Scn5a*-M1875T^+/-^ hearts. Scale bars represent 10 μm. **c)** Quantification of WGA-stained atrial area and atrial cell diameters from transverse sections as depicted in b). Neither parameter was affected by the point mutation (WGA-stained atrial area quantified from LA appendage: N=6 hearts and 65/67 individual ROI per group; LA posterior wall N=6 hearts and 26/29 ROI per group. Atrial cell diameter quantified from LA appendage: N=6 hearts and 14563/13368 individual cells per group; LA posterior wall and 8283/8384 individual cells per group). Data are presented as mean ± SD.

**Figure 3.**
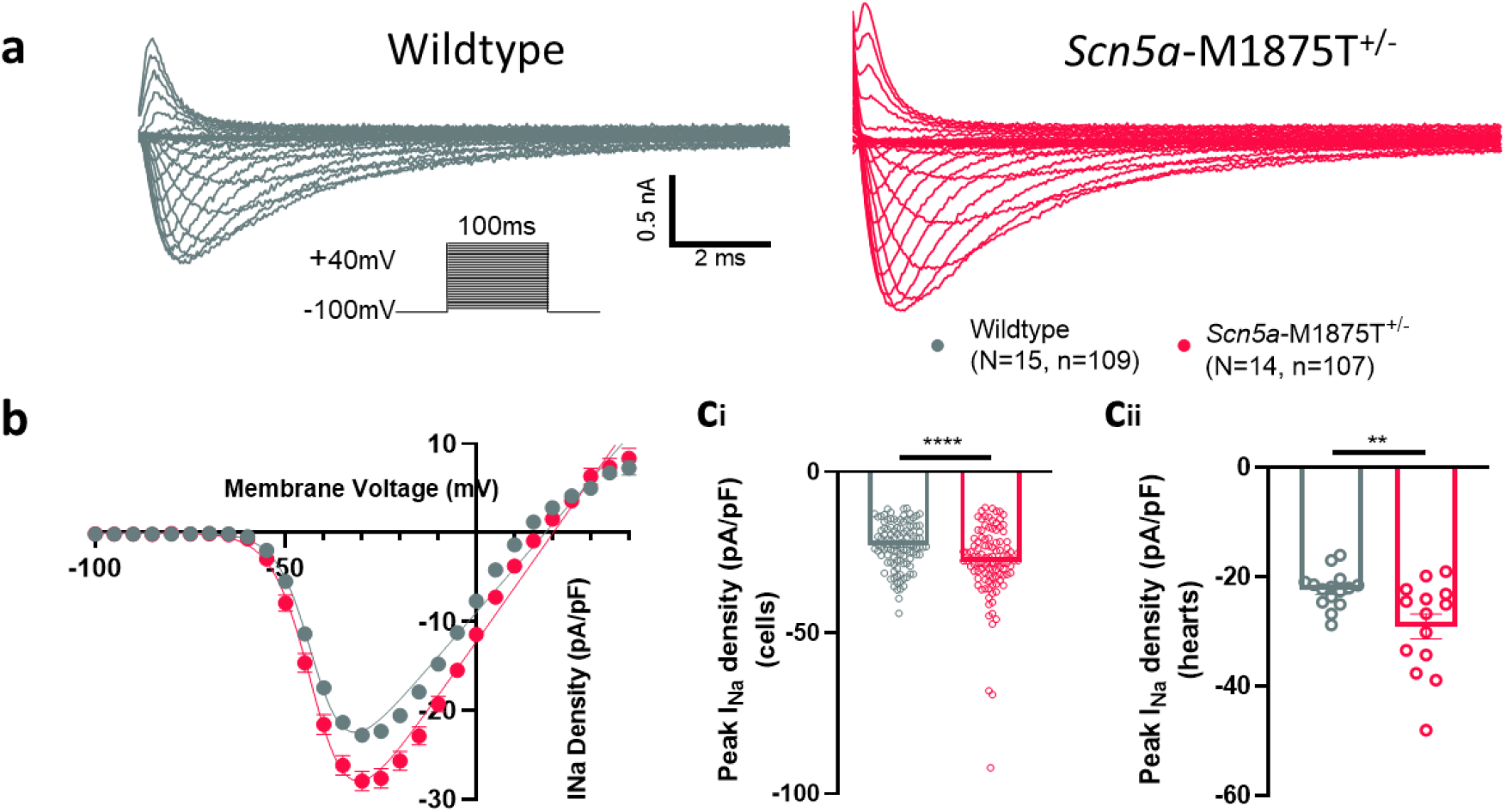
Isolated atrial cardiomyocytes with the *Scn5a*-M1875T^+/-^ mutation have a larger sodium current than wildtypes when measured with whole-cell patch clamp electrophysiology. **a)** Representative sodium current (I_Na_) traces from whole-cell voltage clamp recording of isolated left atrial (LA) cardiomyocytes from wildtype (WT) (grey) and *Scn5a*-M1875T^+/-^ (red) hearts. **b)** Normalised grouped data revealed that the *Scn5a*-M1875T^+/-^ mutation increases peak I_Na_ in LA cardiomyocytes over test potentials ranging from −100 to +40 mV. **c)** At the peak I_Na_ test potential of −30 mV, *Scn5a*-M1875T^+/-^ cardiomyocytes had a significantly larger I_Na_ than WTs, both when comparing individual cells (n=109 WT, n=107 *Scn5a*-M1875T^+/-^) **(ci)** and hearts (N=15 WT, N=14 *Scn5a*-M1875T^+/-^) **(cii)**. Data are presented as the mean ± SEM.

## 3 Results

### 3.1 Viable heterozygous *Scn5a*-M1875T^+/-^ mice show normal cardiac size, structure and basic function

The point mutation previously identified in patients with early familial AF was successfully introduced to exon 28 of the mouse *Scn5a* gene via homologous recombination of a targeting vector. The vector contained the T-C point mutation (CRISPR/cas9-mediated) resulting in methionine-threonine exchange in the Na_v_1.5 protein (Figure 1, Figure S1a). Offspring from both WT x heterozygote (*=Scn5a*-M1875T^+/-^) and heterozygote x heterozygote pairings were viable. No homozygous *Scn5a*-M1875T^-/-^ offspring were born (Figure S1b), suggesting embryonic lethality, as previously reported for other *Scn5a* mutations ^23^. Accordingly, the ratio of WT and heterozygous animals shifted from 1:2 (expected) to approximately 1:3 when heterozygous animals were crossed (Figure S1b). The ratio of male:female sex in offspring approximated 1:1 as expected (Figure S1c).

Age-matched young adult WT and *Scn5a*-M1875T^+/-^ mice displayed similar heart rate, PR-, QRS- and QT-interval regardless of genotype in electrocardiograms recorded awake (Table S1) and during sedation (Table S2). Echocardiography and histological examination excluded overt differences in structure (Table S3, Figure 2a). Neither atrial extracellular matrix content nor atrial cardiomyocyte cell diameter were affected by the mutation (Figure 2b and c, Figure S2a). Accordingly, pro atrial natriuretic peptide protein expression (proANP) was detected in right atria as expected but was not elevated in ventricles (Figure S2b).

### 3.2 Atrial *Scn5a*-M1875T^+/-^ cardiomyocytes have an augmented peak sodium current density

To determine the impact of the M1875T point mutation on peak I_Na_ amplitude and Na_v_1.5 channel gating properties, left atrial cardiomyocytes from WT and *Scn5a*-M1875T^+/-^ mice (8-13 weeks) were isolated and whole-cell patch clamp recordings were performed.

The M1875T variant increased I_Na_ over test potentials ranging from −100 to +40 mV (Figure 3a and b). At a peak test potential of −30 mV, mean left atrial cardiomyocyte I_Na_ density was higher in *Scn5a*-M1875T^+/-^ (−28.0 ± 1.1 pA/pF, n=107 cells) than in WT littermates (−22.8 ± 0.7 pA/pF, n=109 cells, p<0.0001, Figure 3ci). The elevation in I_Na_ was also apparent when recordings were grouped by heart (*Scn5a*-M1875T^+/-^ −29.1 ± 2.2 pA/pF, N=14 vs WT −22.3 ± 0.9 pA/pF, N=15, p=0.0075, Figure 3cii) and after applying hierarchical analysis (*Scn5a*-M1875T^+/-^ −29.1 ± 1.1 pA/pF, n=107 cells, N=14 mice vs WT −22.3 ± 0.9 pA/pF, n=109 cells, N=15 mice, p=0.0093).

Capacitance measurements were not different between genotypes, indicative of similar cell size of atrial cardiomyocytes (Figure S3a).

Whole-cell I_Na_ voltage-dependent inactivation and time-dependent recovery kinetics were not altered in *Scn5a*-M1875T^+/-^ cardiomyocytes compared to WT (Figure S3b and c). Na_v_1.5 expression in hearts of *Scn5a*-M1875T^+/-^ mice at the mRNA and the protein level revealed no difference at either the whole cell or isolated membrane fraction level (Figure S4).

### 3.3 Action potentials from *Scn5a*-M1875T^+/-^ atria have a larger amplitude and faster peak upstroke velocity

Action potentials (APs) were measured in whole left atrial tissue isolated from WT and *Scn5a*-M1875T^+/-^ mice (9-13 weeks) using sharp microelectrodes. Representative AP traces are shown in Figure 4 during stimulation at 100 ms (Figure 4c) and 1000 ms pacing cycle length (PCL) (Figure 4f).

**Figure 4.**
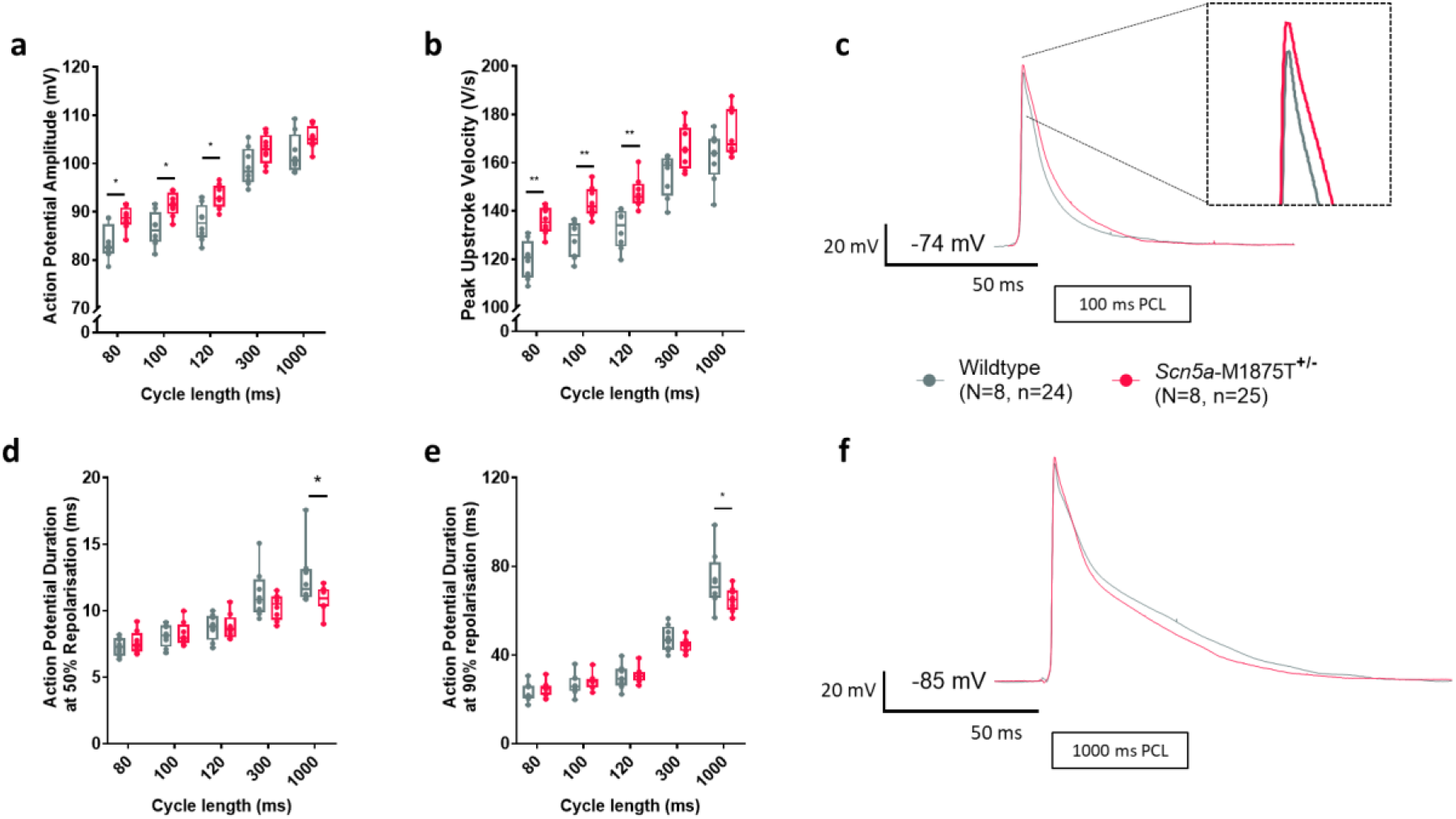
Action potentials from atria with the *Scn5a*-M1875T^+/-^ mutation have a larger action potential amplitude and a faster peak upstroke velocity when measured with the sharp microelectrode technique. **a)** *Scn5a*-M1875T^+/-^ left atria (LA) had a significantly larger action potential (AP) amplitude at shorter pacing cycle lengths (PCLs) of 80-120 ms (p<0.05) and **b)** a significantly faster peak upstroke velocity (dV/dt). **d-f)** When paced at 1000 ms PCL, *Scn5a*-M1875T^+/-^ LA had a significantly shorter AP duration (APD) when measured at 50% (APD50) **(d)** and 90% (APD90) **(e)** repolarisation. *p<0.05, **p<0.01, wildtype (WT) vs *Scn5a*-M1875T^+/-^, N=8 per group, n=24 WT, n=25 *Scn5a*-M1875T^+/-^; ANOVA statistics. Also shown are representative AP traces from WT (grey) and *Scn5a*-M1875T^+/-^ (red) LA when stimulated at 100 ms **(c)** and 1000 ms PCL **(f)**. Data are presented as the mean ± SEM.

Action potential amplitude was significantly larger in *Scn5a*-M1875T^+/-^ murine left atria at all PCLs tested and this effect was more pronounced at shorter cycle lengths (N=8, n=24-25, Figure 4a). The variant resulted in a faster peak upstroke velocity (dV/dt), especially at the shorter cycle lengths (100 ms PCL: WT 128.0 ± 3.3, n=24; *Scn5a*-M1875T^+/-^ 142.8 ± 4.0 mV/ms, n=25, p=0.0282, Figure 4b).

The resting membrane potential ^24^ was not different between genotypes (100 ms PCL: WT −72.4 ± 0.6; *Scn5a*-M1875T^+/-^ −73.1 ± 0.6 mV). Atrial activation times were also similar (100 ms PCL: WT 4.9 ± 0.2; *Scn5a*-M1875T+/- 4.9 ± 0.2 ms) (Table S4). Only at the long PCL of 1000 ms, the AP duration (APD) at 50 and 90% repolarisation was shorter in *Scn5a*-M1875T^+/-^ left atria than in WT (Figure 4d and e), while the APD at 30 and 70% repolarisation was not significantly different (Table S4). Similarly, optical mapping data showed no APD differences at PCLs tested.

### 3.4 Flecainide-induced atrial conduction slowing and post-repolarisation refractoriness is less pronounced in *Scn5a*-M1875T^+/-^ atria

Optical mapping of WT and *Scn5a*-M1875T^+/-^ whole left atrial tissue was performed to test effects of the heterozygous *Scn5a*-M1875T mutation on atrial conduction. Left atria were superfused with the open channel sodium channel blocker flecainide (1 μM, clinically used concentration) to determine the response of *Scn5a*-M1875T^+/-^ and WT left atria. While conduction velocity was unchanged at baseline, flecainide slowed conduction less in *Scn5a*-M1875T^+/-^left atria (conduction velocity difference at 100 ms PCL −6 ± 1 cm/s, n=12, p=0.0357) than in WTs (conduction velocity difference at 100 ms PCL −10 ± 1 cm/s, n=12, p=0.0357, Figure 5a and b). We also investigated flecainide-induced changes in left atrial refractoriness as it is known that flecainide induces post-repolarisation refractoriness using rapid atrial pacing. Representative optical AP recordings show 1:1 capture in the *Scn5a*-M1875T^+/-^ left atria with flecainide, while several stimuli in the WT left atria did not elicit APs (Figure 5c). All time-controlled left atria (no flecainide, same experimental duration) were successfully paced with 1:1 capture down to 80 ms PCL. Loss of 1:1 capture began at longer PCLs in WT (7/12 atria lost 1:1 capture at PCLs ≤ 120 ms, Figure 5d) compared to *Scn5a*-M1875T^+/-^ (1/12 atria lost 1:1 capture at PCLs ≤ 120 ms, p=0.0185) left atria. The diastolic pacing threshold remained consistent throughout experiments. Thus, flecainide induced less pronounced post-repolarisation refractoriness in the *Scn5a*-M1875T^+/-^ left atria.

**Figure 5.**
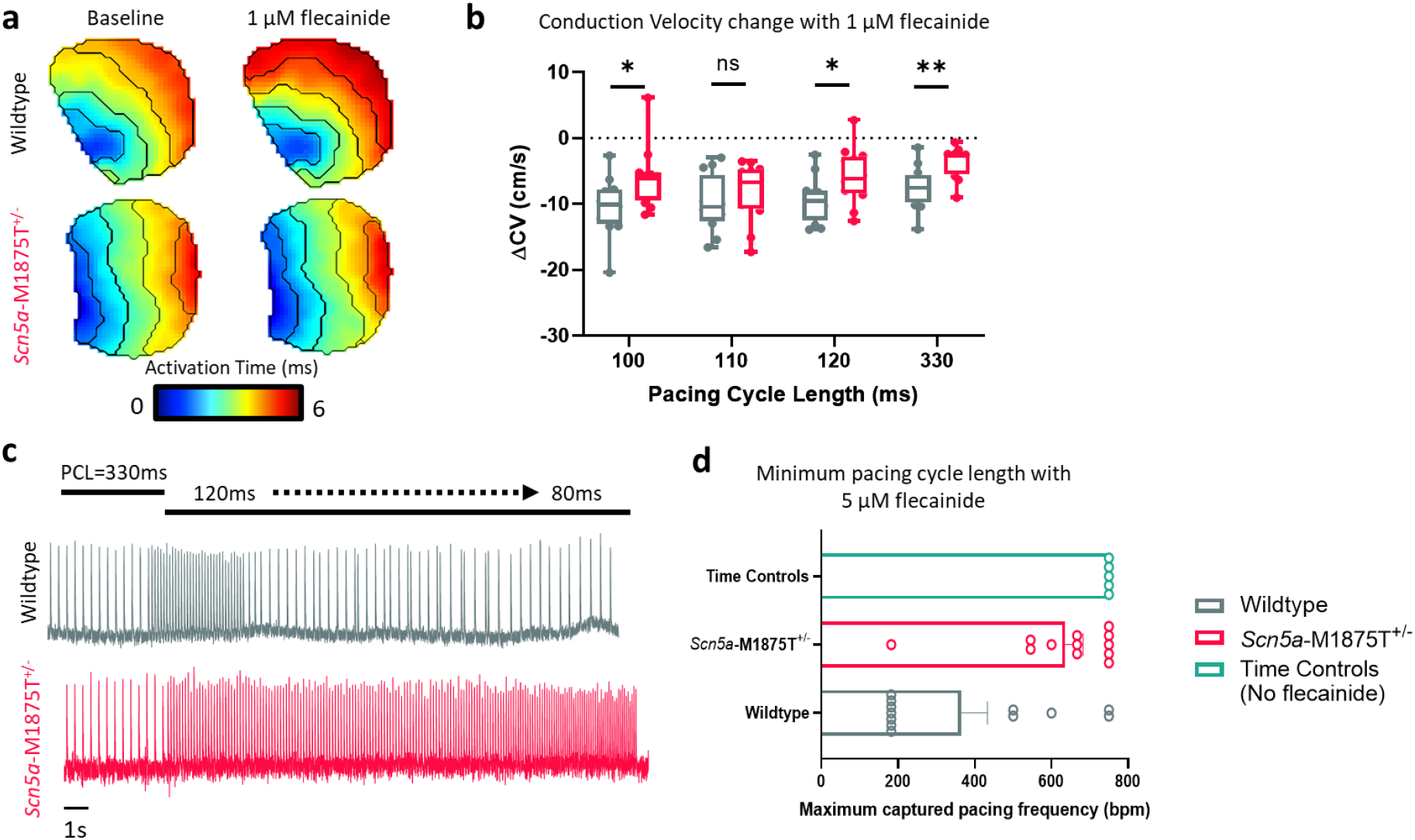
The response to the anti-arrhythmic agent flecainide is reduced in *Scn5a*-M1875T^+/-^ atria in optical mapping. **a)** Example left atrial (LA) activation maps from wildtype (WT, top panels) and *Scn5a*-M1875T^+/-^ (bottom panels) mice. Left panels show activation maps at baseline, right panels show activation of the same atria following exposure to 1 μM flecainide for 20 minutes. **b)** Grouped data showing change in conduction velocity (ΔCV) following exposure to 1 μM flecainide in WT (grey) and *Scn5a*-M1875T^+/-^ (red) LA for 20 mins. **c)** Example traces of optical action potentials (APs) recorded following further treatment of WT (top, grey) and *Scn5a*-M1875T^+/-^ (bottom, red) LA with 5 μM flecainide for 15 mins. **d)** Grouped data showing minimum pacing cycle length (PCL) at which 1:1 stimulus capture was maintained in WT (grey) and *Scn5a*-M1875T^+/-^ (red) LA following exposure to 5 μM flecainide. Time control data (green) shows minimum PCL at which 1:1 stimulus capture was maintained in atria that were not exposed to flecainide but had been under experiment conditions for the same time period (35 mins from baseline recording). N=12 per group at 330 ms-100 ms PCL. Atria were excluded from further analysis at shorter PCLs if 1:1 capture was lost, only data in steady state was used. N=5 for time control experiments. *p<0.05, **p<0.01, WT vs *Scn5a*-M1875T^+/-^; ANOVA statistics. Data are represented as the mean ± SEM.

## 4 Discussion

### 4.1 Main findings

Our study describes the effects of the familial atrial fibrillation (AF) mutation *Scn5a*-M1875T^+/-^ in a newly generated murine model. Key effects are that the M1875T *Scn5a* mutation leads to an increased atrial action potential upstroke velocity and amplitude, and a selective increase in the early cardiac sodium current (I_Na_). Atrial cardiomyocyte capacitance and size, as well as cardiac size and function, are preserved. The effect of the sodium channel blocker flecainide is dampened in *Scn5a*-M1875T^+/-^ atria. Our measurements in this new murine model confirm that a selective increase of I_Na_ can cause familial AF as observed in the family affected, and suggest that commonly used concentrations of sodium channel blockers may be less effective in familial forms of AF with a selective increase in I_Na_ than in other types.

### 4.2 Gain-of-function properties of *Scn5a*-M1875T^+/-^ sodium channels in the murine atrium

Our data show a gain-of-function variant, namely an increased early sodium current as evidenced by an augmented action potential amplitude and upstroke velocity, and larger I_Na_ in the *Scn5a*-M1875T^+/-^ variant.

In contrast to findings in HEK293 cells ^13^, there was no depolarising shift of Na_v_1.5 channel inactivation in murine left atrial cardiomyocytes observed. Instead, we show a similarly augmented I_Na_ without alterations in channel gating properties. Generation of the murine mutant model allowed us to study mutated sodium channels in cardiac tissue within the presence of the greater protein complex including α- and β-subunits and other membrane proteins, a complete cardiomyocyte contractile apparatus and all other cardiac cell types. Others have reported similar differences between HEK293 and cardiac model systems to the differences reported here ^25^.

Cardiac sodium channels have been observed to form dimers ^26^ and different regions of the channel protein have been implicated in dimerization. Structural analysis following the hypothesis of Na_v_1.5 dimerization via C-terminal interaction reveals that the surface of residue Met1875 of one Na_v_1.5 cytosolic C-terminus will interact with Ala1924 of a second Na_v_1.5 ^27^. This prediction suggests that the location site containing the mutation in our model could provide a structural basis for altered Na_v_1.5-Na_v_1.5 channel interaction further to be investigated.

It is unlikely that the M1875T mutation increases the *late* sodium current (I_Na,l_), as APs we measured were not prolonged by the mutation, in line with the initial report of the mutation in HEK cells ^13^. A lack of AP prolongation clearly differs to findings in the gain-of-function mutation Δ*KPQ-Scn5a^+/-^* mutant murine model which shows an increase in I_Na,l_^17, 28^ with prolonged atrial and ventricular AP duration, especially at longer pacing cycle lengths. This suggests that the M1875T mutation acts differently to, and is distinct from, *SCN5A* gain-of-function mutations leading to prolonged repolarisation and long QT syndrome, in concordance with the broad spectrum of phenotypic outcomes resulting from mutations in the same ion channel gene ^29^.

An enhanced sodium influx into cardiomyocytes should alter the sodium-calcium homeostasis ^30, 31^, modifying intracellular sodium and calcium concentrations and signalling in cardiomyocyte microdomains ^32, 33^. This study found normal heart and atrial cardiomyocyte size. Nonetheless, chronic sodium overload can result in structural changes in the heart with ageing. In the future, the M1875T mouse model may be used to study long-term effects of a selective increase in the early I_Na_ with ageing.

### 4.3 Effect of *Scn5a*-M1875T^+/-^ on conduction and refractoriness

While structural alterations of the atria contribute to atrial conduction disturbances ^34^, defects in the cardiac sodium channel can also cause conduction defects leading to AF ^11, 12, 35^. Unlike other *SCN5A* mutations ^35^, this variant shows a dampened response to flecainide both on conduction and post-repolarisation refractoriness ^36^. Flecainide is a clinically used sodium channel blocker that inhibits cardiac I_Na_ via blocking the pores of open Na_v_1.5 channels ^37^. The differential response to flecainide is likely due to the increased early sodium influx through the mutated Na_v_1.5 channels, enabling preservation of conduction and activation properties in atrial cardiomyocytes when sodium channels are inhibited, leading to an enhanced activation reserve.

Murine models have limitations due to differences between mice and human, but due to consistency in sodium currents have been useful in characterising sodium channel mutations ^17, 23, 38^. Identified mutation-induced changes at the cellular and organ level in this model appear sufficient to explain AF in the original family. Studies in human cardiomyocytes ^31^ and in atrial engineered heart tissue would be desirable to assess the effect of the M1875T mutation in human models in the future.

### 4.4 Conclusion

The *Scn5a*-M1875T^+/-^ variant causes a selective increase in the early cardiac sodium current, leading to an increased activation reserve and reduced refractoriness, whilst structure and contractile function are preserved. These findings can explain the familial occurrence of atrial ectopy and atrial fibrillation in the absence of reported severe heart disease. More widely they suggest a mechanism by which altered cardiomyocyte sodium current can predispose to atrial fibrillation. Our data also show that the M1875T gain-of-function mutation decreases the effectiveness of sodium channel blockers such as flecainide, which may have implications for treatment.

## Data Availability Statement

The datasets generated during the current study are available from the corresponding author on reasonable request.

## Author Contribution Statement

LF and PK designed the research; BVS generated the mouse model in TRAM upon request by PK and LF;

MO, LCS, CO, SBS, SA, JSR, SNK, AM, DD, LFo, LF and APH conducted experiments, analysed data and performed statistical analysis;

KG, DP, APH, PK, LF supervised experiments and analysis.

MO, LCS, CO, SBS and LF wrote the manuscript together with all co-authors. All co-authors critically reviewed the manuscript.

## Acknowledgments

We thank Hartwig Wieboldt, Clara Apicella and Olivia Grech for technical assistance and acknowledge the UKE Microscopy Imaging Facility (UMIF) for expert support.

