## Supplement for "Increased early sodium current causes familial atrial fibrillation and dampens effect of flecainide"

### Supplemental Figures

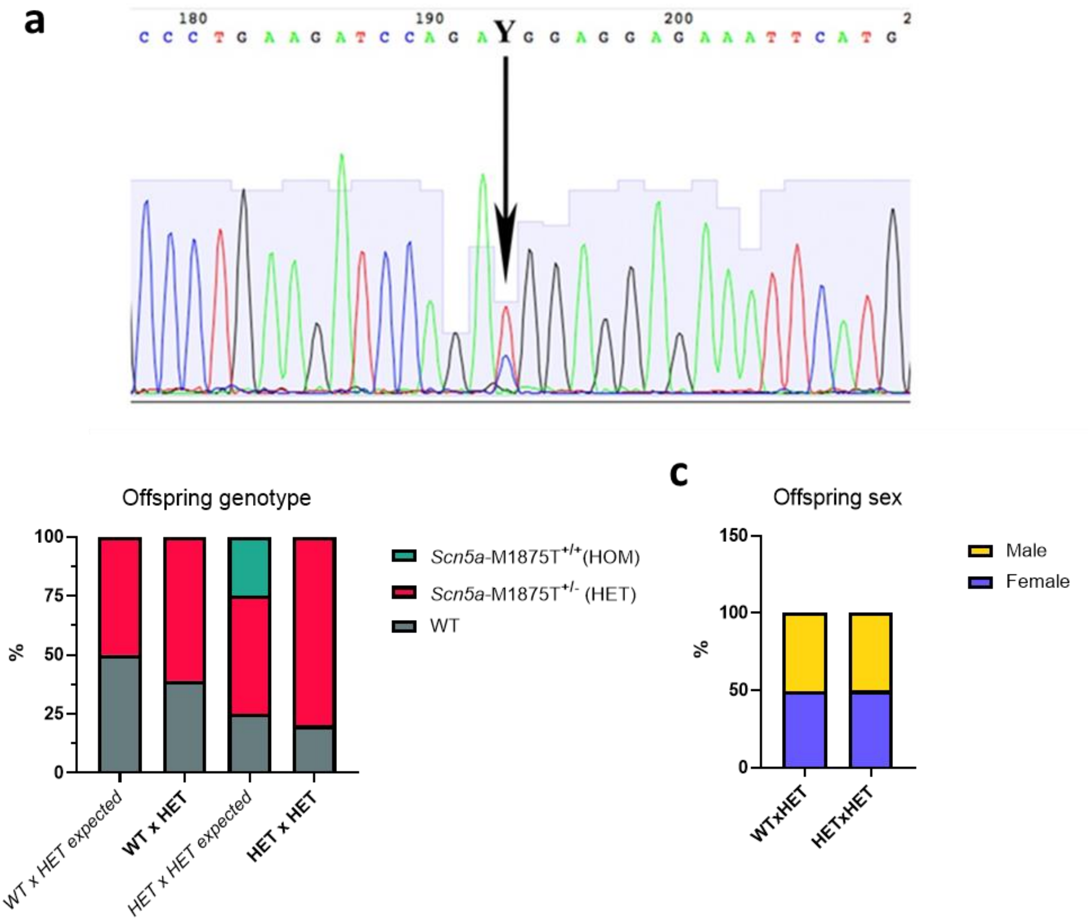


**Figure S1: Sequencing analysis of F1 offspring, and subsequent offspring genotype and sex ratios. a)** Sequencing analysis of the heterozygous F1 offspring animal number 8 (Figure 1b). Position of the T-C mutation, which corresponds to M1875(7)T amino acid exchange in the protein sequence, is labeled by an arrow. **b)** Observed offspring genotype and **c)** sex distribution, analysed from 90 litters (901 animals in total), pairing with WTxHET or HETxHET animals. WT, wildtype; HET, heterozygous; HOM, homozygous.

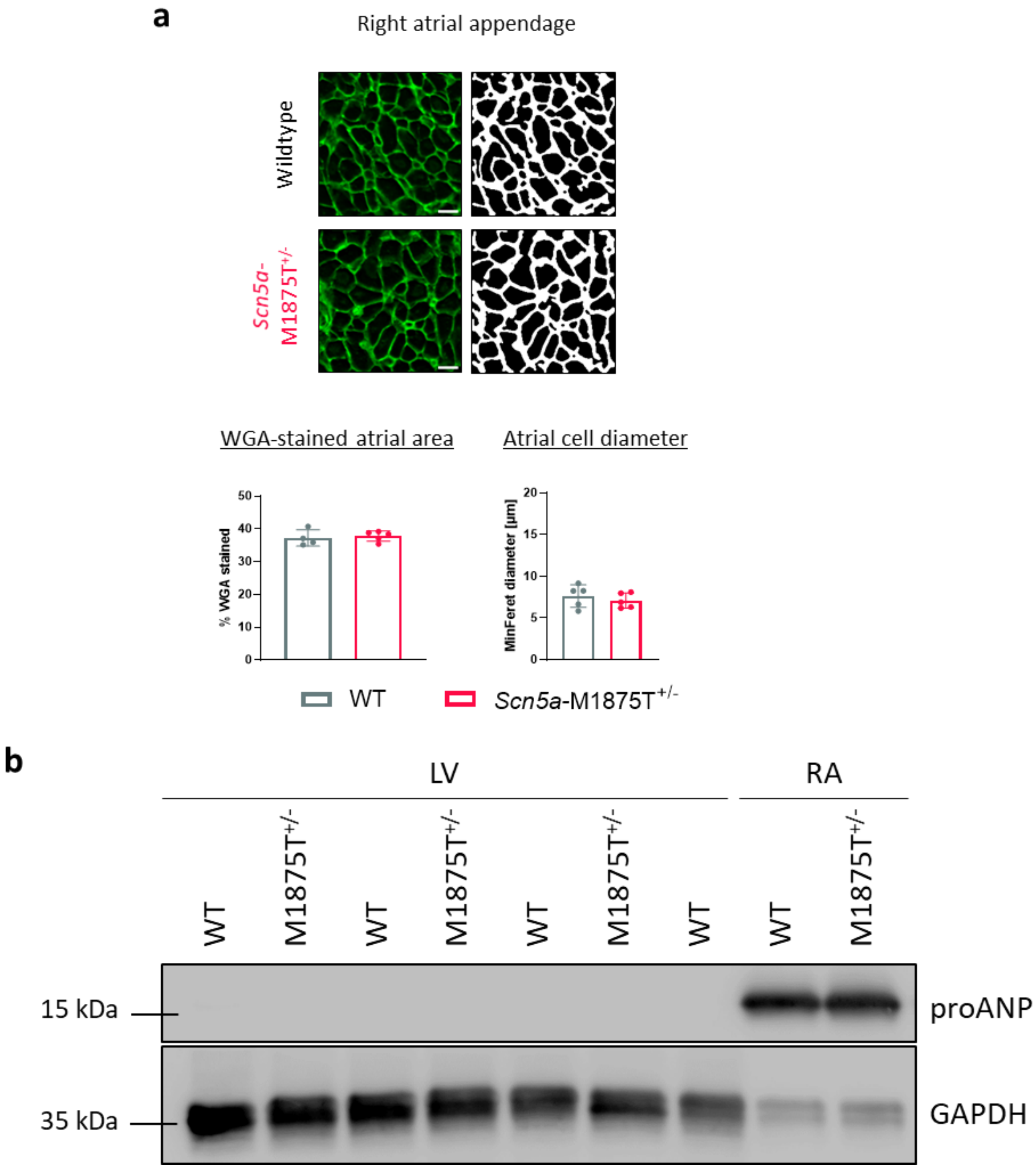


**Figure S2: Histological phenotyping of wildtype and *Scn5a*-M1875T$^{+/-}$ right atria and ventriclular and right atrial proANP protein level assessment**

**a)**

Exemplary immunofluorescence images from atrial regions of interest (ROI) of wheat germ agglutinin (WGA, green) staining and corresponding binary images for quantification in left atrial (LA) appendage and LA posterior wall from wildtype (WT, left) and *Scn5a*-M1875T$^{+/-}$ (right) mice. Scale bars represent 10 µm. Graphs show quantification of WGA-stained atrial area and atrial cell diameters from transverse sections as depicted above. Neither parameter was affected by the point mutation at young adult age (WGA-stained atrial area quantified from LA appendage: N=6 hearts and 22/21 individual ROI per group; Atrial cell diameter quantified from N=6 hearts and 8548/11883 individual cells per group. Data are presented as mean ± SD. **b)** Immunoblot probing for pro-ANP in left ventricular (LV) as well as right atrial (RA) tissue. GAPDH expression serves as loading reference with reduced loading for the atrial samples. There was no ventricular signal for proANP, excluding overt heart failure.

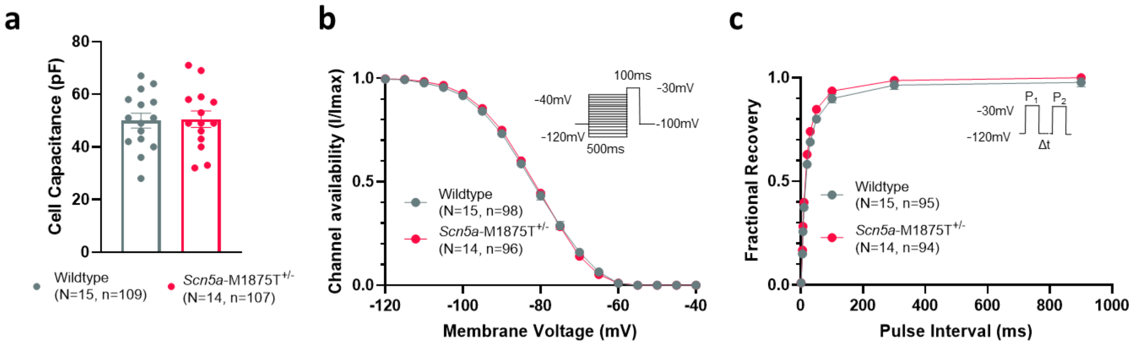


**Figure S3: Measurements of cell capacitance, voltage-dependent inactivation and time-dependent recovery kinetics of $I_{Na}$ using patch clamp electrophysiology. a)** Cell capacitance measurements exclude overt hypertrophy, **b)** voltage-dependent inactivation and **c)** time-dependent recovery kinetics of $I_{Na}$ in isolated left atrial cardiomyocytes from wildtype and *Scn5a*-M1875T$^{+/-}$ mice.

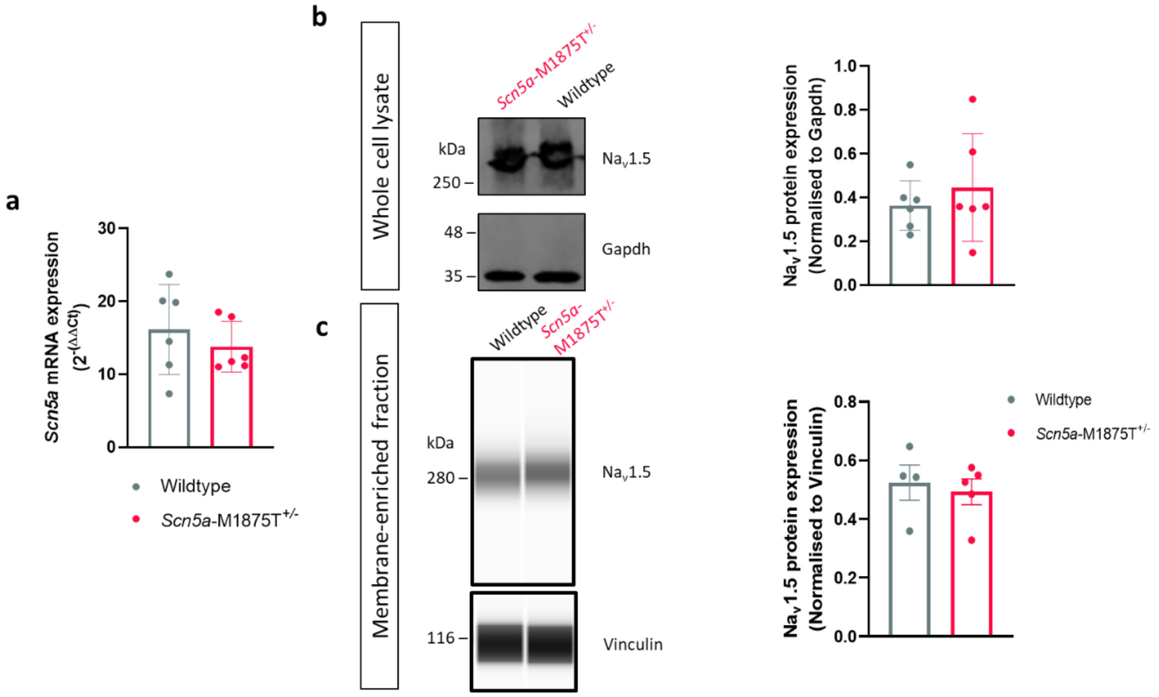


**Figure S4: Quantification of *Scn5a* mRNA and $Na_v$1.5 protein expression in the *Scn5a*-M1875T$^{+/-}$ and wildtype mouse hearts, when measured at the whole cell and isolated membrane fraction level. a)** *Scn5a* mRNA quantification of LA tissue samples from *Scn5a*-M1875T$^{+/-}$ as well as WT mice measured using RT-qPCR. Expression levels were normalised against GAPDH (n=6 per group).

**b)** $Na_v$1.5 protein expression levels in left atrial (LA) tissue samples from *Scn5a*-M1875T$^{+/-}$ and wildtype mice were measured using western blotting. GAPDH was used as a loading control. A representative blot is shown (left) with band quantification (right) (n=6 per group). **c)** $Na_v$1.5 protein quantification in membrane-enriched fractions of LA tissue samples from *Scn5a*-M1875T$^{+/-}$ (n=4) and WT (n=5) mice using capillary-based automated western blotting. Vinculin was used as a loading control. The detected protein amount is visualised (left) and area under peak was quantified (right). There were no obvious changes between genotypes.

### Supplemental Tables

| Parameters | Wildtype | *Scn5a*-M1875T$^{+/-}$ | P value |
|---|---|---|---|
| **n** | 19 | 19 | NA |
| **Age (weeks)** | 16 ± 0.5<br>(12-18) | 15 ± 0.5<br>(12-18) | 0.12 |
| **Heart rate (beats/min)** | 681 ± 7.0<br>(610-727) | 691 ± 6.3<br>(655-755) | 0.30 |
| **RR interval (ms)** | 88 ± 1.0<br>(83-99) | 87 ± 0.8<br>(80-92) | 0.28 |
| **P duration (ms)** | 13 ± 0.4<br>(9.8-16.5) | 13 ± 0.6<br>(7.3-18.0) | 0.23 |
| **PR duration (ms)** | 30 ± 0.7<br>(26-36) | 30 ± 0.6<br>(26-36) | 0.57 |
| **QRS duration (ms)** | 14 ± 0.3<br>(12-17) | 13 ± 0.2<br>(12-15) | 0.28 |
| **QT duration (ms)** | 46 ± 1.1<br>(40-53) | 47 ± 1.0<br>(39-53) | 0.55 |
| **R amplitude (mV)** | 0.35 ± 0.04<br>(0.10-0.76) | 0.37 ± 0.04<br>(0.11-0.78) | 0.74 |

**Table S1: Electrocardiogram (ECG) characteristics of conscious wildtype and *Scn5a*-M1875T$^{+/-}$ mice.** Data are expressed as mean ± SEM (range).

| **Parameters** | **Wildtype** | ***Scn5a*-M1875T$^{+/-}$** | **P value** |
|---|---|---|---|
| **n** | 23 | 21 | NA |
| **Age (weeks)** | 14 ± 0.7<br>(9-20) | 14 ± 0.6<br>(10-20) | 0.95 |
| **Heart rate (beats/min)** | 433 ± 8.2<br>(388-494) | 423 ± 5.3<br>(393-467) | 0.30 |
| **RR interval (ms)** | 139 ± 2.7<br>(110-155) | 142 ± 1.8<br>(129-153) | 0.26 |
| **P duration (ms)** | 16 ± 1.2<br>(10-25) | 15 ± 1.3<br>(11-25) | 0.70 |
| **PR duration (ms)** | 38 ± 1.1<br>(24-47) | 38 ± 1.3<br>(24-49) | 0.83 |
| **QRS duration (ms)** | 15 ± 0.8<br>(10-23) | 16 ± 0.9<br>(12-25) | 0.88 |

**Table S2: Electrocardiogram (ECG) characteristics of unconscious wildtype and *Scn5a*-M1875T$^{+/-}$ mice.** Data are expressed as mean ± SEM (range).

| Parameter | Wildtype | *Scn5a*-M1875T+/- | P value |
|---|---|---|---|
| n | 21 | 26 | N/A |
| Age (weeks) | 15 ± 0.6<br>(9-18) | 14 ± 0.6<br>(9-20) | 0.30 |
| Body weight (g) | 28 ± 1.2<br>(19-42) | 28 ± 0.8<br>(23-38) | 0.84 |
| Heart rate (beats/min) | 438 ± 7.5<br>(382-501) | 427 ± 6.4<br>(380-499) | 0.29 |
| LA diameter (mm) | 2.2 ± 0.1<br>(1.6-2.7) | 2.3 ± 0.1<br>(1.6-2.7) | 0.35 |
| LA area ($mm^2$) | 9.7 ± 0.3<br>(7.1-12.7) | 9.4 ± 0.3<br>(6.6-11.9) | 0.54 |
| IVS, d (mm) | 0.8 ± 0.03<br>(0.5-1.1) | 0.9 ± 0.02<br>(0.7-1.1) | 0.16 |
| IVS, s (mm) | 1.1 ± 0.05<br>(0.7-1.5) | 1.1 ± 0.04<br>(0.8-1.5) | 0.46 |
| LVID, d (mm) | 3.9 ± 0.1<br>(3.0-4.7) | 3.9 ± 0.1<br>(3.2-4.4) | 0.88 |
| LVID, s (mm) | 2.9 ± 0.1<br>(1.7-3.6) | 2.8 ± 0.1<br>(1.7-3.5) | 0.62 |
| LVPW, d (mm) | 0.9 ± 0.04<br>(0.6-1.2) | 0.9 ± 0.04<br>(0.7-1.3) | 0.57 |
| LVPW, s (mm) | 1.1 ± 0.1<br>(0.8-1.6) | 1.2 ± 0.1<br>(0.8-1.5) | 0.44 |
| LVEF (%) | 52 ± 3.0<br>(29-82) | 53 ± 2.2<br>(39-83) | 0.62 |
| LVFS (%) | 25 ± 2.0<br>(10-49) | 28 ± 1.6<br>(19-51) | 0.34 |
| LV mass (mg) | 97 ± 5.0<br>(66-137) | 104 ± 4.0<br>(70-144) | 0.32 |
| LV mass/BW | 3.5 ± 0.1<br>(2.7-4.6) | 3.7 ± 0.1<br>(2.8-4.8) | 0.16 |
| RVID, d (mm) | 1.7 ± 0.07<br>(1.3-2.2) | 1.7 ± 0.04<br>(1.4-1.9) | 0.82 |

**Table S3: Echocardiographic measurements of wildtype and *Scn5a*-M1875T+/- mice.** Values are mean ± SEM (range). Mann-Whitney test used for statistical analysis; IVS, interventricular septum; LVID, left ventricular internal diameter; LVPW, left ventricular posterior wall; LVEF, left ventricular ejection fraction; LVFS, left ventricular fractional shortening; LV mass, left ventricular mass; BW, body weight: LA diameter, left atrial diameter; LA area, left atrial area; RVID, right ventricle internal diameter; d, diastole; s, systole.

| | Wildtype | | | | | *Scn5a*-M1875T+/- | | | | |
|---|---|---|---|---|---|---|---|---|---|---|
| | Pacing cycle length (ms) | | | | | Pacing cycle length (ms) | | | | |
| | **1000.00** | **300.00** | **120.00** | **100.00** | **80.00** | **1000.00** | **300.00** | **120.00** | **100.00** | **80.00** |
| **APD30** | 6.3 ±0.2 | 6.5 ±0.2 | 5.5 ±0.1 | 5.3 ±0.1 | 4.8 ±0.1 | 5.7 ±0.1 | 6.0 ±0.1 | 5.6 ±0.1 | 5.3 ±0.1 | 5.0 ±0.1 |
| **APD50** | 12.5 ±0.5 | 11.3 ±0.4 | 8.8 ±0.2 | 8.1 ±0.2 | 7.3 ±0.1 | 10.8* ±0.2 | 10.3 ±0.2 | 8.8 ±0.2 | 8.3 ±0.2 | 7.6 ±0.2 |
| **APD70** | 38 ±1.9 | 24 ±0.9 | 15 ±0.5 | 13 ±0.4 | 12 ±0.3 | 31 ±0.8 | 21 ±0.4 | 15 ±0.4 | 14 ±0.4 | 12 ±0.3 |
| **APD90** | 73 ±2.6 | 47 ±1.2 | 30 ±1.1 | 27 ±1.0 | 23 ±0.8 | 65* ±1.1 | 44 ±0.6 | 31 ±0.7 | 28 ±0.8 | 25 ±0.7 |
| **APA (mV)** | 102 ±0.8 | 99 ±0.8 | 88 ±0.7 | 86 ±0.7 | 84 ±0.7 | 105 ±0.5 | 103 ±0.6 | 93* ±0.5 | 91* ±0.5 | 88* ±0.5 |
| **dV/dt (mV/ms)** | 162 ±2.1 | 154 ±1.8 | 132 ±1.6 | 128 ±1.5 | 120 ±1.6 | 172 ±2.0 | 166 ±1.8 | 147** ±1.3 | 144** ±1.3 | 135** ±1.1 |
| **RMP (mV)** | -84 ±0.3 | -82 ±0.3 | -73 ±0.4 | -72 ±0.4 | -71 ±0.5 | -84 ±0.5 | -82 ±0.5 | -75 ±0.5 | -73 ±0.4 | -72 ±0.3 |
| **AT (ms)** | 4.6 ± 0.1 | 4.7 ± 0.1 | 4.7 ± 0.1 | 4.8 ± 0.1 | 5.1 ± 0.2 | 4.6 ± 0.2 | 4.7 ± 0.2 | 4.7 ± 0.1 | 4.8 ± 0.1 | 5.1 ± 0.1 |

**Table S4: Action potential characteristics in wildtype and *Scn5a*-M1875T$^{+/-}$ left atria at all pacing cycle lengths tested using the sharp microelectrode technique.** Stated are means ± SEM for: action potential durations (APDs) at 30 (APD30), 50 (APD50), 70 (APD70) and 90% (APD90) repolarisation, action potential amplitude (APA), peak upstroke velocity (dV/dt), resting membrane potential (RMP) and activation times (ATs), at pacing cycle lengths (PCLs) of 80, 100, 120, 300 and 1000 ms. At 1000 ms PCL, APD50 and APD90 were significantly reduced in *Scn5a*-M1875T$^{+/-}$ left atria relative to wildtype. APA was significantly larger in *Scn5a*-M1875T$^{+/-}$ mutants at PCLs of 80-300 ms. dV/dt was faster in mutants when tested at PCLs of 80-120 ms. No significant changes in RMP or AT were detected at any PCL. Values are shown here for reference and as mean ± SEM. *p<0.05, **p<0.01, WT vs *Scn5a*-M1875T$^{+/-}$, N=8 per group, n=24 WT, n=25 *Scn5a*-M1875T$^{+/-}$).

### Supplemental Methods

#### 1 Generation of the *Scn5a*-M1875T construct and knock-in mouse model

##### 1.1 Targeting construct design:

The 3.7 kb genomic region, containing exon 28 of the *Scn5a* gene and right flanking region (RFR) was PCR amplified from mouse genomic DNA and (T-C) mutated using combination of subsequent PCR reactions with help of oligonucleotide pairs SCN5aA1d / SCN5aA1r, SCN5aA2d / SCN5aA2r and SCN5aMUT, subcloned and sequenced. The 5.1 kb left flanking region (LFR) containing exons 25-27, and intronic sequences was PCR amplified from mouse genomic DNA using oligonucleotides SCN5aBcld / SCN5aBclr and subcloned. The 1.3 kb genomic region, containing exon 24 and intronic sequences was added to the LFR. All individual clones were verified by sequencing and assembled into the final targeting construct (pSCN5a_targ3) in the order depicted in Figure 1a. The pBluescript based plasmid backbone together with the negative selection marker (thymidine kinase cassette) were added to the left flanking region (not shown). The 1.9 kb DNA fragment, containing a positive selection marker (neomycin cassette flanked by two LoxP sites) was cloned between LFR and RFR.

##### 1.2 Southern blot DNA probes cloning:

The 1.4 kb HR probe was PCR amplified from mouse genomic DNA using oligonucleotides SCN5aHRd2 and SCN5aHRr2, subcloned and sequenced. The So1, So2, So3, So4 and So5 probes were PCR amplified from mouse genomic DNA, subcloned and sequenced using pairs of oligonucleotides: SCN_SO1D/SCN_SO1R, SCN_SO2D/SCN_SO2R, SCN_SO3D/SCN_SO3R, SCN_SO4D/SCN_SO4R, and SCN_SO5D/SCN_SO5R accordingly.

##### 1.3 Scn5a exon 28 specific gRNA's selection and cloning for the CRISPR/cas9 system:

Three gRNA targets were selected in a close proximity to the mutation site. These gRNAs were cloned in plasmid gRNA_Cloning Vector (gift from George Church (Addgene plasmid # 41824))[1] digested with the *Afl*II restriction endonuclease, using the Gibson assembly methods with help of oligonucleotide pairs: SCN5a_ Insert_F/SCN5a_ Insert_R, SCN5a_ Insert_F2/SCN5a_ Insert_R2, and SCN5a_Insert_F3/SCN5a_Insert_R3 resulting in plasmids pgRNA_SCN1, pgRNA_SCN2, and pgRNA_SCN3 accordingly.

**1.4 ES cell transfection and selection of targeted clones**

CV19 ES cells (passage 13 [129Sv x C57BL/6J]) were expanded in HEPES-buffered Dulbecco's modified Eagle's medium supplemented with 15% fetal bovine serum (PAA), nonessential amino acids, L-glutamine, β-mercaptoethanol, 1,000 U per mL of recombinant leukemia inhibitory factor (LIF) (MERCK Millipore), and antibiotics (penicillin [100 U/ml] and streptomycin [100 μg/ml]). For electroporation, 2 x $10^7$ cells were resuspended in 0.8 ml Capecchi buffer (20 mM HEPES [pH 7.4], 173 mM NaCl, 5 mM KCl, 0.7 mM $Na_2HPO_4$, 6 mM dextrose, 0.1 mM β-mercaptoethanol)[2]. The targeting vector DNA pSCN5a_targ3 (100 μg) was electroporated together with 70 μg of each plasmid DNA's: pgRNA_SCN1, pgRNA_SCN2, pgRNA_SCN3 and hCAS9 (cas9 coding plasmid was a gift from George Church (Addgene plasmid # 41815))[1] at 25 μF and 400V in 0.8 mm electroporation cuvettes (Gene Pulser; Bio-Rad). After electroporation, cells were cultivated for 10 min at room temperature and plated onto ten 100-mm diameter culture dishes containing a gamma-irradiated monolayer of mouse primary G418-resistant fibroblast feeder cells. Thirty-two hours later, 350 μg of G418 (Invitrogen) per mL and 0.2 μM 2'-deoxy-2'-fluoro-β-D-arabinofuranosyl-5-iodouracil (FIAU) (Moravek Biochemicals and Radiochemicals, USA) were added to the culture medium. The medium was replaced every day, and colonies were picked and analysed 8 days after plating.

**1.5 DNA Southern blot analysis**

Positively targeted ES cell clones were analyzed using the Southern-blot DNA method. Approximately 5-10 μg of genomic DNA was digested with *Eco*RI, fractionated on 0.8% agarose gels, and transferred to GeneScreen nylon membranes (NEN DuPont). The membranes were hybridized with a $^{32}P$-labeled 1.4 kb HR probe and washed with (final concentrations) 0.5x SSPE (1x SSPE is 0.18 M NaCl, 10 mM $NaH_2PO_4$, and 1mM EDTA [pH 7.7]) and 0.5% sodium dodecyl sulfate at 65°C. After first screening, correctly targeted clones were proven using the *Bam*HI and additionally *Hind*III digestion, using a $^{32}$P-labeled 1.8 kb probe So3 containing internal sequences to the targeted homology. The Southern blot analysis of DNA samples isolated from F1 mouse tail biopsy using probe So3 is presented on (Figure 1b).

**1.6 Blastocyst injection**

Correctly targeted ES cells from 9C clone were injected into 3.5-day B6D2F1 blastocysts. Routinely, we are injecting 12 to 14 ES cells into one blastocoele. After injection, blastocysts were kept in KSOM medium and subsequently transferred into the uteri of 2.5-day pseudopregnant CD-1 foster mice. The mice carried pups to term. Chimeras were identified by their agouti coat colour contribution. For the germ-line transmission high percentage male chimaeras were crossed to the C57BL/6J female mice. Heterozygous agouti offsprings (*Scn5a*-M1875$^{+/-}$) were confirmed by Southern blot analysis (Figure 1b), and the mutated part of exon 28 was PCR amplified using primers pair SCN5a_Sequenc_F/ SCN5a_Sequenc_R and sequenced (Figure 1c and Supplementary Figure 1a). Mice were kept in specific pathogen-free animal facilities.

**1.7 Deletion of the neo cassette**

The deletion of the Neo cassette (Figure 1a) was performed by crossing mice with the total CRE deleter transgenic animals and verified by PCR analysis and further confirmed by the Southern blot analysis (not shown), using different probes, including the probe containing the Neo cassette sequences.

**1.8 List of oligonucleotides used in the study:**

| | |
|---|---|
| SCN5aA1d | *TGATATC*GTGAGGAGCTGGAAGCCTTGAG |
| SCN5aA1r | CCGTCTGGATCTTCAGGGCATCCATCTC |
| SCN5aA2d | TGAAGATCCAGACGGAGGAGAAATTCATGG |
| SCN5aA2r | *TGCGGCCGC*ATTTATACACGAAGCTTAGTGAGAAGTG |
| SCN5aMUT | GAGATGGATGCCCTGAAGATCCAGACGGAGGAGAAATTCATGG |
| SCN5aBcld | *TGTCGAC*TGCATGCACTTGATGGCCTCAC |

| | |
|---|---|
| SCN5aBclr | *TGAATTC*CCTAGTCCATCTCCTCCTAACA |
| SCN5aHRd2 | GACTAAGTGATGCTAAAACACAT |
| SCN5aHRr2 | TGTGTGTATGTGTAGAGGTTGTC |
| SCN_SO1D | *CAGTCTCAAGCGTCTCTTGGAGTCGA*CCTCACTACAGCCGCACACTCAC |
| SCN_SO1R | *CTGCTCTAGACGTCTCTGAGAGTCGA*CCAAGAAAGTCAGGCTAGCAGAGAG |
| SCN_SO2D | *CAGTCTCAAGCGTCTCTTGGAGTCGAC*AGAATAAATAACATCTACTCCATAAG |
| SCN_SO2R | *CTGCTCTAGACGTCTCTGAGAGTCGAC*AAGTCAAGGAAGACATAGCAG |
| SCN_SO3D | *CAGTCTCAAGCGTCTCTTGGAGTCGA*CTAGGCTGATGCAGTGCTGAAG |
| SCN_SO3R | *CTGCTCTAGACGTCTCTGAGAGTCGAC*ACATGTACAGTGTGTTTCCCGTT |
| SCN_SO4D | *CAGTCTCAAGCGTCTCTTGGAGTCGAC*TCCTGCTGGCTTTTGATTGTGC |
| SCN_SO4R | *CTGCTCTAGACGTCTCTGAGAGTCGAC*AAAAGGGACATCTCTTGGGAAACT |
| SCN_SO5D | *CAGTCTCAAGCGTCTCTTGGAGTCGA*CAGAAAACTTGAGCCAATCCAC |
| SCN_SO5R | *CTGCTCTAGACGTCTCTGAGAGTCGA*CCCCGTGGGCACCGTTTAG |
| SCN5a_ Insert_F | TTTCTTGGCTTTATATATCTTGTGGAAAGGACGAAACACCGGCCCTGAAGATCCAGATGG |
| SCN5a_ Insert_R | GACTAGCCTTATTTTAACTTGCTATTTCTAGCTCTAAAACCCATCTGGATCTTCAGGGCC |
| SCN5a _Insert_F2 | TTTCTTGGCTTTATATATCTTGTGGAAAGGACGAAACACCGCTCGGGGAGTCTGGGGAGA |
| SCN5a _Insert_R2 | GACTAGCCTTATTTTAACTTGCTATTTCTAGCTCTAAAACTCTCCCCAGACTCCCCGAGC |
| SCN5a _Insert_F3 | TTTCTTGGCTTTATATATCTTGTGGAAAGGACGAAACACCGTAGGAGATCTTGGAAGGAT |
| SCN5a _Insert_R3 | GACTAGCCTTATTTTAACTTGCTATTTCTAGCTCTAAAACATCCTTCCAAGATCTCCTAC |
| SCN5a_Sequenc_F | GCCCTGTCCGACTTTGCCGAT |
| SCN5a_Sequenc_R | CTGGCGGAAGAGGAAGGAAGCAT |

Sequences used for cloning are italicized. Scn5a-M1875T mutated nucleotide is underlined.

#### 2 mRNA and protein expression levels

##### 2.1 RNA preparation and quantitative real-time PCR

RNA was purified from LA and left ventricle tissue as previously described [3]. Briefly, LA and left ventricle tissue was homogenised using the Precellys homogeniser (Precellys) and the RNA was purified using the RNEasy mini kit (Qiagen). Total RNA (1 µg) was used as a template for reverse transcription with the High-Capacity cDNA Reverse Transcription kit (Thermo Fisher) according to the manufacturer's protocol. A TaqMan® probe to *Scn5a* (Mm01342518_m1; Applied Biosystems) was used to quantify mRNA levels by RT-PCR using TaqMan™ Universal PCR Master Mix. *Gapdh* (Mm99999915_g1; Applied Biosystems) was used for normalisation. The results of six separate experiments, each calculated from three technical replicates, were pooled using the ΔΔCt method.

##### 2.2 Western blotting methods

Western blotting of whole LA, RA and LV protein lysate was carried out as previously described [3]. Briefly, snap-frozen cardiac chambers were lysed in ice-cold radioimmune precipitation buffer (Sigma) with Halt Protease and Phosphatase Inhibitor mixture (Thermo Fisher) using a mechanical Precellys 24 tissue homogeniser (Precellys) or cryogrinding. Total protein concentration was quantified using the DC Protein Assay kit (Bio-Rad). After normalisation of protein concentration using SDS reducing sample buffer, samples were denatured at 95°C for 5 minutes before being resolved by SDS-PAGE and immunoblotted using primary antibodies directed against Nav1.5 (D9J7S; Cell Signaling) or pro-ANP (ab180649, Abcam). Quantification of GAPDH protein expression (14C10; Cell Signaling) served as a loading control. Protein-antibody complexes were visualised using HRP-based chemiluminescence.

To collect a membrane enriched LA protein lysate, the LA was homogenised in ice cold lysis buffer (50 mM Tris-Base (pH 7.5), 2 mM EDTA, 5 mM EGTA, 5 mM DTT, 0.05% Digitonin) supplemented with Halt Protease and Phosphatase Inhibitor mixture. Cytoplasmic fraction was collected through centrifugation at 17,000 g for 30 minutes at 4°C. The resulting pellet was resuspended in ice cold lysis buffer supplemented with 1% Triton-X before repeating centrifugation. The resulting supernatant was defined as a membrane-enriched fraction. Protein concentration was determined using DC protein assay (Bio-Rad). $Na_v1.5$ protein expression was determined in the membrane fraction using Western capillary electrophoresis (WES) method (ProteinSimple, San Jose, CA). Left atrial membrane fractions from wild-type mice and mice possessing the mutation were loaded into WES 13-well plates for separation using capillary electrophoresis (66-440 kDa) following manufacturer's instructions. Antibodies directed against $Na_v1.5$ (Cell Signalling) and Vinculin (Cell signalling) were both used at a dilution of 1:50 in antibody diluent (ProteinSimple). The relative amount of each protein was analysed through the areas under peaks from the chemiluminescence chromatograms by Compass for SW software (ProteinSimple).
